# Accessory-cell-free differentiation of hematopoietic stem and progenitor cells into mature red blood cells

**DOI:** 10.1101/2022.09.14.507311

**Authors:** Yelena Boccacci, Nellie Dumont, Yannick Doyon, Josée Laganière

## Abstract

The culture and *ex vivo* engineering of red blood cells (RBCs) can help characterize genetic variants, model diseases, and may eventually spur the development of applications in transfusion medicine. In the last decade, improvements to the *in vitro* production of RBCs have enabled efficient erythroid progenitor proliferation and high enucleation levels from several sources of hematopoietic stem and progenitor cells (HSPCs). Despite these advances, there remains a need for refining the terminal step of *in vitro* human erythropoiesis — i.e., the terminal maturation of reticulocytes into erythrocytes — so that it can occur without feeder or accessory cells and animal components. Here, we describe the near-complete erythroid differentiation of cultured RBCs (cRBCs) from adult HSPCs in accessory-cell-free and animal-component-free conditions. The approach improves post-enucleation cell integrity and cell survival, and enables subsequent storage of cRBCs for up to 42 days in classical nutritive solution conditions, without any specialized equipment. We foresee that these improvements will facilitate the characterization of RBCs derived from gene-edited HSPCs.

**KEY POINTS:**

- Erythroid progenitors were differentiated into fully mature RBCs in a medium free of accessory cells
- Cultured RBCs can be stored up to 42 days in a standard nutritive solution

## INTRODUCTION

Erythropoiesis is a multistep process in which hematopoietic stem and progenitor cells (HSPCs) differentiate into red blood cells (RBCs). HSPCs first differentiate into immature, enucleated reticulocytes in the bone marrow. Reticulocytes subsequently complete their maturation into erythrocytes in the peripheral circulation, where they lose RNA, organelles (e.g., mitochondria), and various cell surface markers (e.g., the transferrin receptor CD71, the integrin-associated protein CD47, and the glycoprotein IV CD36).^1–5^ The resulting mature erythrocytes are thus smaller and denser than reticulocytes and feature increased hemoglobin-carrying and deformability potential.^3,4,6^

Enormous efforts have been devoted to reproducing erythropoiesis *in vitro* for various applications, such as the production of blood (for transfusion) and laboratory reagents (for serologic tests).^7–17^ These *in vitro* methods are also useful to characterize genome-edited erythroid cells differentiated from HSPCs.^18–26^

Cultured RBCs (cRBCs), an umbrella term that designates *in-vitro*-generated red cell products, can be produced from a number of cell sources with varying efficacy.^15,27^ Cell lines and induced pluripotent stem cells are promising, unlimited cell sources for *in vitro* blood production, but enucleation and subsequent erythrocyte maturation are much less efficient with these sources.^28–31^ Thus, HSPCs remain a widely used cell source in the laboratory.^7–14,32–37^

While current protocols starting from HSPCs yield high proliferation of progenitor cells and high enucleation rates, they rarely enable reticulocytes to fully mature into RBCs without feeder or accessory cells — i.e., the final maturation step remains incomplete.^7–14,32–37^ In addition, post-enucleation cell survival during maturation is rarely reported. Therefore, a protocol that fully reproduces reticulocyte maturation and that is compatible with genome editing could be, for instance, an additional tool to complement the development of autologous HSPC-derived gene therapies.^38^

We thus aimed to identify conditions that would support erythroid expansion and differentiation, and maximize cell maturation to obtain cRBCs that would resemble circulating RBCs. We describe Erythroid Differentiation Medium for Terminal Maturation (EDM-TM) — a four-phase culture system to differentiate HSPCs into RBCs that is free of accessory and feeder cells. EDM-TM achieves robust proliferation of erythroid progenitors, high enucleation rates, and a near-complete maturation of reticulocytes into erythrocytes. EDM-TM also enhances post-enucleation red cell integrity and maintenance, as well as storage survival at 4 °C.

## MATERIALS AND METHODS

### Cell culture

EDM-TM is an optimized version of EDM — a 3-phase, erythroid differentiation culture protocol for the *in vitro* production of RBCs from HSPCs.^10^ The composition of the medium, with the concentrations of additives used in each culture phase, is shown in **Table 1,** and details on the additives (i.e., vendor, catalog number) can be found in **Table S2**. Two versions were developed: EDM-TM-1 (i.e., the original version) and EDM-TM-2 (i.e., a version optimized to reduce the costs of EDM-TM-1).

**Table 1:** Culture media composition.

| Additives |  | EDM<sup>10</sup> |  |  |
| --- | --- | --- | --- | --- |
| Basal medium | Glutamine | 2 mM |  |  |
|  | Insulin | 10 ug/mL |  |  |
|  | Heparine | 2 U/mL |  |  |
|  | Human plasma | 5% |  |  |
| Phase 1<br>Day 0-8<br>1.10<sup>4</sup> cells/mL | Hydrocortisone | 0.001 mM |  |  |
|  | IL-3 | 5 ng/mL |  |  |
|  | SCF | 100 ng/mL |  |  |
|  | EPO | 3 U/mL |  |  |
|  | Transferrin | 330 ug/mL |  |  |
| Phase 2<br>Day 8-12<br>1.10<sup>5</sup> cells/mL | SCF | 100 ng/mL |  |  |
|  | EPO | 3 U/mL |  |  |
|  | Transferrin | 330 ug/mL |  |  |
| Phase 3<br>Day 12-19<br>1-5.10<sup>6</sup> cells/mL |  | **EDM** | **EDM-TM-1** | **EDM-TM-2** |
|  | EPO | 3 U/mL | 3 U/mL | 3 U/mL |
|  | FeIII-EDTA | - | - | 4uM |
|  | Transferrin | 330 ug/mL (low) | 1 mg/mL (high) | 330 ug/mL (low) |
|  | Human plasma | 5% | 20% (*Day 15) | 20% (*Day 15) |
|  | Slow agitation | - | Day 19 | Day 19 |
|  | Filtration | Day 19 | Day 22 | Day 22 |
| Phase 4<br>Day 19-32<br>5.10<sup>6</sup> cells/mL | EPO | 3 U/mL | 3 U/mL | Remove on day 22 |
|  | FeIII-EDTA | - | - | 4uM |
|  | Transferrin | 330 ug/mL (low) | 1 mg/mL (high) | 330 ug/mL (low) |
|  | Plasma | 5% | 20% | 20% |
|  | Slow agitation | - | + | + |

In both versions, G-CSF-mobilized peripheral blood stem cells (mPBSCs) from distinct donors (**Table S1**, purchased from AllCells [Alameda, CA, USA]) were incubated at 37°C/5% CO_2_ in a humid atmosphere throughout the culture. The basal medium consisted of Iscove’s Modified Dulbecco’s Medium supplemented with 2 mM glutamine, 10 μg/mL human insulin, 330 μg/mL human transferrin, 2 U/mL human heparin, and 5% human plasma that was ABO-compatible with HSPC donor cells. In *phase 1*, mPBSCs were thawed and seeded at 1.10^4^ cells/mL in basal medium supplemented with 0.001 mM hydrocortisone, 5 ng/mL human recombinant interleukin 3 (IL-3), 100 ng/mL human recombinant stem cell factor (SCF), and 3 U/mL human recombinant erythropoietin (EPO). On day 4, cells were diluted 1/5 in the same medium (**Table 1**). In *phase 2* (initiated on day 8), cells were enumerated with a Nucleocounter NC-250 (Chemometec, Somerville, MA, USA) and diluted at 1.10^5^ cells/mL in medium supplemented with 100 ng/mL SCF and 3 U/mL EPO (**Table 1**). In *phase 3* (initiated on day 12), EDM-TM-1 included the following modifications relative to the original EDM protocol: transferrin concentration was increased to 1 mg/mL starting on day 12, and plasma concentration was increased to 20% starting on day 15 (**Table 1; Figure 1A**). In *phase 4* (initiated on day 19), cells were gently agitated and were otherwise maintained in the same conditions as in Phase 3. On day 22 (timing of maximal enucleation), cells were filtered with Acrodisc WBC Syringe Filter 25 mm (VWR, Mont-Royal, QC, Canada) to isolate enucleated cells, which were then cultured in phase 4 conditions. The maturation of enucleated reticulocytes was continued up until day 32 (**Table 1; Figure 1A**). For storage at 4 °C, AS-3 storage solution was diluted 1/2 with the culture medium.

**Figure 1:**
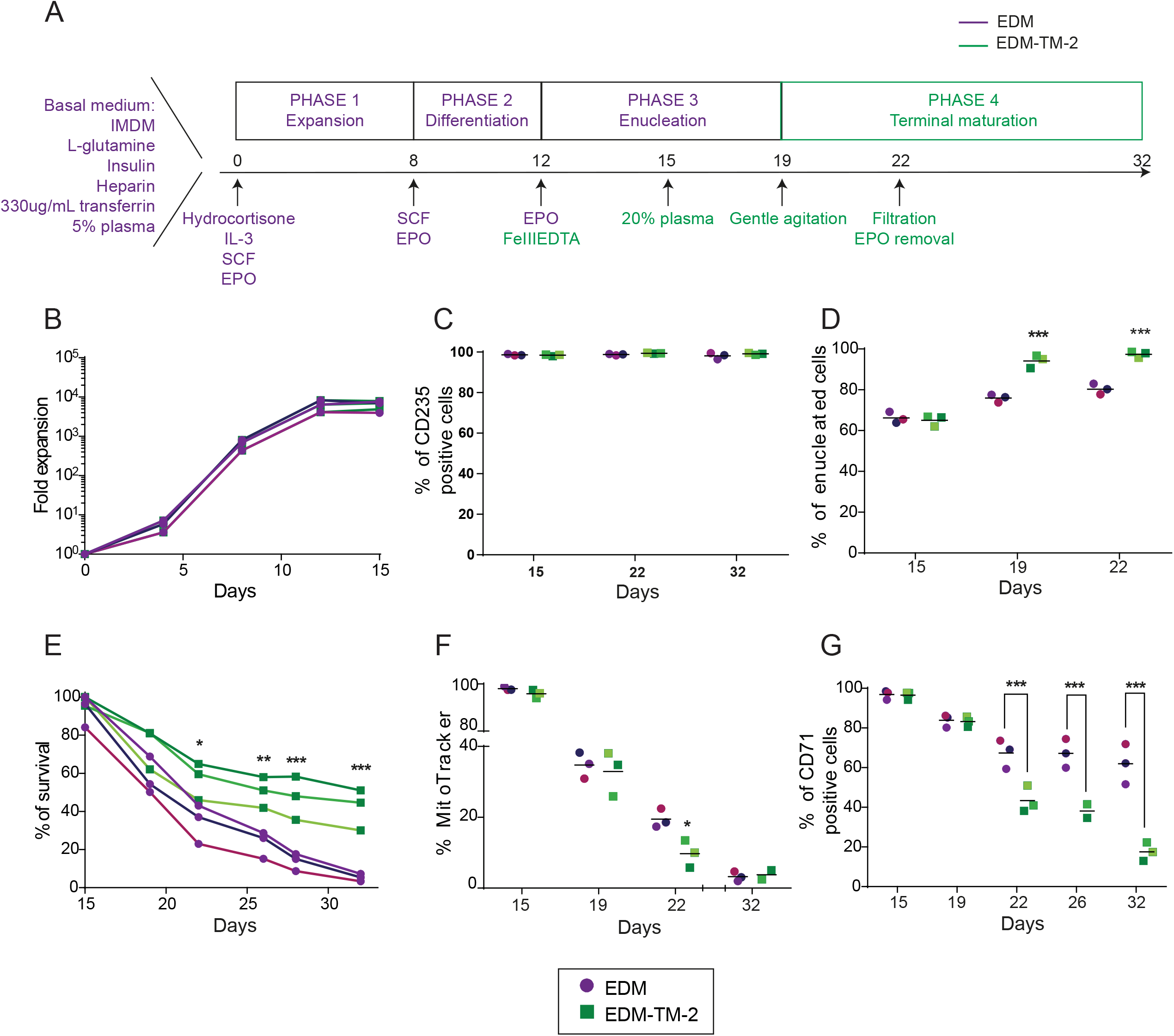
Efficient production and maintenance of mature cRBCs with EDM-TM-2. (A) Schematic overview of EDM-TM-2, with modifications relative to EDM indicated in green. (B) Erythroid expansion of HSPCs from day 0 to day 15. HSPCs from three distinct donors are illustrated with different shades of the same color. (C) Percentage of CD235a+ (late erythroid differentiation marker and RBC marker) cells; (D) Percentage of enucleated, SYTO-Green+ cells on days 15, 19 and 22; (E) Percentage of cRBC survival expressed as the percentage of the maximum number of cells obtained through the culture; (F) Percentage of MitoTracker+ (mitochondria marker) cells; (G) RBC maturation expressed as the decreasing percentage of CD71+ cells. *P <0.05; **P <0.01; ***P <0.001

In EDM-TM-2, the following changes were implemented to reduce costs: (1) 1 mg/ml transferrin was either replaced with 4 μM FeIII-EDTA from day 12 up until end of culture, or reduced to 330 μg/ml from day 22 up until end of culture; (2) EPO was removed from day 22 up until end of culture. The other steps were the same as those described above for EDM-TM-1 (**Table 1; Figure 2A**).

**Figure 2:**
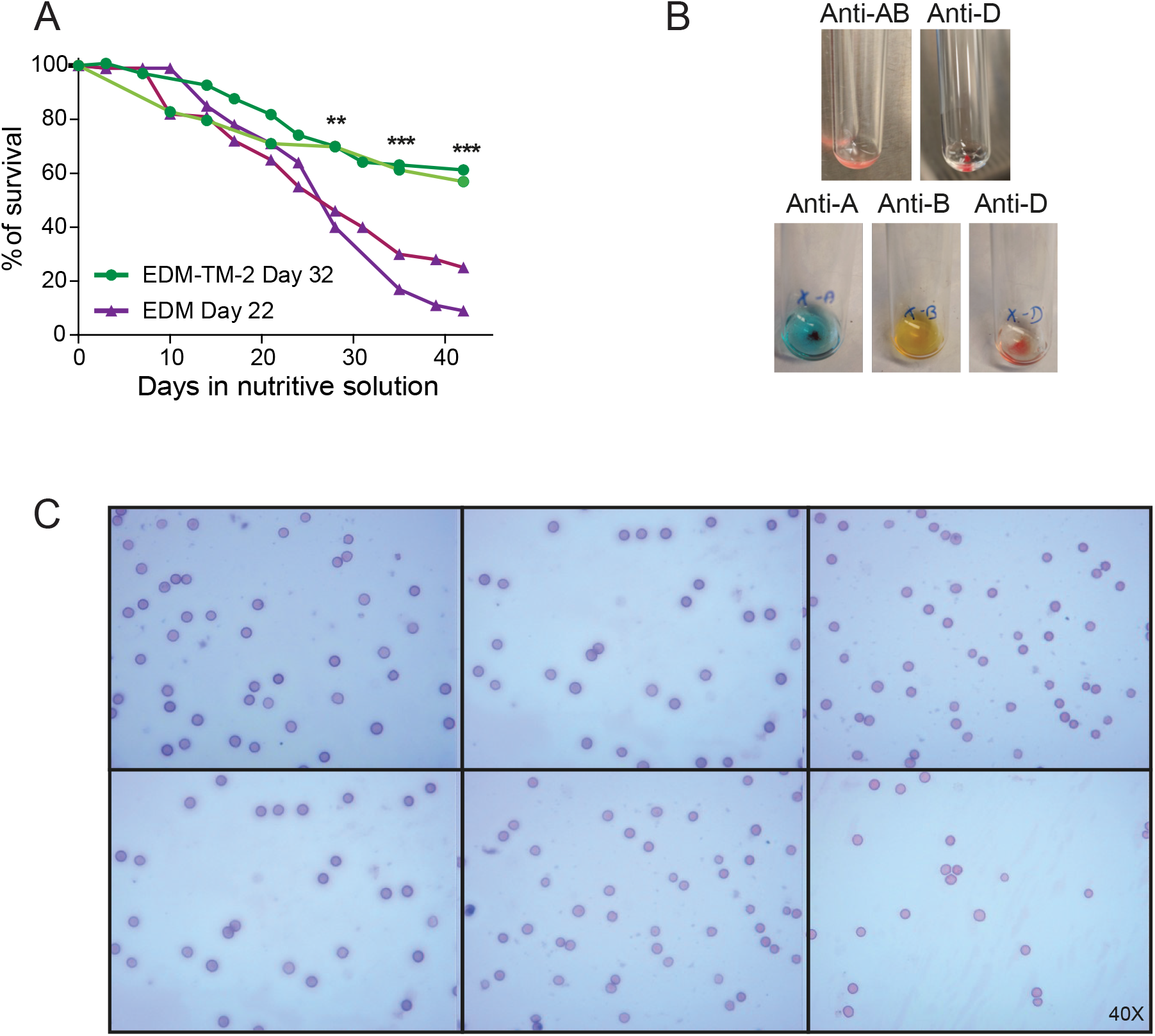
EDM-TM-2-derived cRBCs exhibit enhanced long-term storage. (A) Conservation of EDM-TM-2-derived cells (cultured up to day 32) and EDM-derived cells (cultured up to day 22) in nutritive solution up to 42 days, expressed as the percentage of the initial number of cells placed in nutritive solution; (B) Images acquired following hemagglutination using anti-AB, anti-A, anti-B and anti-D reagents; (C) Microscopy images of cRBCs acquired on day 42 of storage (day 74 post-protocol initiation) after Wright-Giemsa colorations (40X).

In both versions, cell medium was changed every 3 or 4 days throughout phases 3 and 4 by centrifuging cells at 1200 rpm for 10 minutes. Cell concentration was gradually increased during phases 3 and 4 by resuspending reticulocytes at 1.10^6^ cells/mL on day 12, 3.10^6^ cells/mL on day 15, and 5.10^6^ cells/mL from day 19 up until end of culture.

### Flow cytometry analysis of erythropoiesis and reticulocyte maturation

The following antibodies and dyes were used as markers of erythroid differentiation and maturation for analysis by flow cytometry: anti-CD71-APC (transferrin receptor - erythropoiesis marker; expected to decrease as maturation progresses), anti-CD235a-FITC (glycophorin A - late erythropoiesis marker and RBC marker), anti-CD47-BV421 (integrin-associated protein – reticulocyte maturation marker), anti-CD36-FITC (glycoprotein IV – early erythropoiesis marker; expected to disappear at the reticulocyte stage), 7AAD (viability marker until enucleation), 100 nM SYTO 13 green (DNA marker for nucleated cells), 150 nM MitoTracker™ Orange CMTMRos (mitochondrial loss marker), and 1 mL of RetiCount (i.e., thiazole orange; residual RNA marker) per 5.10^5^ cells.

Fluorescent minus one (FMO) conditions were used to position the boundary for the positive-cell population and gating. Adult RBCs were used as control to confirm that the MitoTracker and RetiCount signals were not due to background or autofluorescence. The Attune NxT Flow Cytometer (ThermoFisher, Waltham, MA, USA) was used, and analysis was performed with FCS express 6 flow research edition. Mean fluorescent intensity (MFI) was obtained with the software to determine the level/intensity of expression of markers.

### Phenotypic characterization

Wright-Giemsa (Sigma-Aldrich, Oakville, ON, Canada) staining was performed following centrifugation with Cytospin3 (ThermoFisher, Waltham, MA, USA) and methanol fixation. Images were acquired using light microscopy at a magnification of 40X. Hemagglutination tests with serological anti-AB, anti-A, anti-B, and anti-D were performed and images were acquired 15 minutes after adding antibodies to 2.10^6^ cells.

For confocal microscopy and 3D reconstructions, cells (1-2.10^5^) were stained 15 minutes at 37 °C with 15 μM Carboxyfluorescein succinimidyl ester and washed once in Hank’s Balanced Salt Solution. Slides were prepared with Cytospin3 and visualized with Quorum Wave FX (Quorum Technologies, Puslinch, ON, Canada). Images were acquired with an ImageEM camera (Hamamatsu, EM-CCD 512×512 pixels) using the Volocity 4 software (Quorum Technologies, Puslinch, ON, Canada).

### Statistical analysis

Data were analyzed with two-way repeated measures analysis of variance (ANOVA) with Dunn-Sidak correction to assess differences between methods at different time points in culture period. Analyses were carried out using Graphpad Prim version 6 (Graphpad Software. CA, USA).

## RESULTS

### Development of EDM-TM-1

EDM was selected as a starting point for optimization because it is an established erythroid differentiation culture protocol for the *in vitro* production of RBCs from adult mobilized HSPCs that provides substantial progenitor expansion, differentiation and enucleation.^10^ This protocol has been used in clinical settings and is widely used in the genome editing field.^18–21^ However, this 3-phase culture protocol neither supports the post-enucleation cell maintenance of reticulocytes nor their maturation into erythrocytes.^10^ Accordingly, we kept the expansion (Phase 1) and differentiation (Phase 2) steps intact but modified the components of the enucleation step (Phase 3) and added a terminal maturation step (Phase 4) (**Figure 1A**).

For this iterative process to complement the EDM system, we tested components identified in various previous studies for their positive impact.^7,9,12,13,32,39–41^ We tested (1) the addition of a higher plasma concentration (20% vs. 5% in EDM) which may improve enucleation, differentiation, and post-enucleation cell integrity *in vitro*;^7,13^ (this plasma addition was started on day 15 as enucleation greatly increases between day 15 and 19), (2) mild agitation, which has been shown to promote maturation^9,32,41^, starting on day 19 since starting earlier leads to increased cell death^41^; and (3) higher transferrin concentrations (1 mg/ml vs. 0.33 mg/ml) in phase 3, which may increase iron intake during the late differentiation of erythroblasts and facilitate the reticulocyte maturation steps.^12,13,32,39,40^ The post-enucleation culture period was also extended with a fourth phase, so that cells spend more time in conditions that promote terminal maturation (summarized in **Figure S1A** and supported by the data in **Supplementary Figures 1 and 2**).

Starting from mPBSC, a mean expansion of 22,000 folds (**Figure S1B**) and >90% of CD235a+ cells (**Figure S1C**) were obtained after 15 days of culture, indicating robust expansion and full erythroid commitment (results from six HSPC donors).Then, EDM-TM-1 consistently yielded >60% enucleation on day 22 (**Figure S1D**).

Both 1 mg/mL transferrin and agitation improved post-enucleation cell maintenance (**Figure S1E**) and better supported RBC maturation compared to EDM, as shown by the decreasing levels of CD71 (**Figure S1F**).

Using a 20% plasma concentration from day 15 improved enucleation levels (**Figure S1G**). Moreover, while only 10%-30% of cells were alive by day 32 with 5% plasma (**Figure S1E** and **S1H**), survival improved with 20% plasma (**Figure S1H**). Taken separately, both agitation (**Figure S1I**) and 1 mg/mL transferrin (**Figure S1J**) also had a positive, additive effect on post-enucleation cell maintenance in 20% plasma conditions.

Agitation from day 19 (**Figure S1L**) and 1 mg/mL transferrin from day 12 (**Figure S1M)** better supported RBC maturation (as measured by decreased levels of CD71), in contrast to 20% plasma which had a limited effect (**Figure S1K**).

The maximum level of enucleation was reached on day 22, which was thus the optimal day for filtration to isolate enucleated RBCs. Together, these changes established EDM-TM-1 (**Table 1**).

### Optimizations to reduce costs in EDM-TM-2

EDM-TM-1 was further optimized to reduce the costs associated with unessential components (**Figure S2A** and **Table 1**). As expected based on the literature, EPO was dispensable post-enucleation (**Figure S2 B-C**) and was therefore removed.^42,43^ Consistent with the reduced importance of transferrin once hemoglobin synthesis is over,^3,4^ using 330 μg/ml (vs. 1 mg/ml) was sufficient for cell maintenance and maturation beyond day 22 (**Figure S2 D-E**). Furthermore, higher transferrin concentrations could be replaced with 4 μM FeIII-EDTA from days 12 to 22 without impairing cell maintenance or maturation post-enucleation (**Figure S2 F-G**), as previously suggested.^31^ Of note, removing FeIII-EDTA on day 22 did not impair cell maintenance (**Figure S2H**) but compromised terminal maturation (**Figure S2I**); it was thus maintained up until the end of the culture. **Figure S2A** summarizes the changes that were retained in EDM-TM-2.

Other variations in media composition were previously shown to help culture RBCs or reduce costs (e.g., removing insulin, replacing transferrin by optiferrin or progressively adding nutritive solution).^13,44^ However, these changes were suboptimal in the present setting and were thus not included in the final protocol (data not shown).

### Improved maturation and maintenance of cRBCs with EDM-TM-2 compared to EDM

To thoroughly characterize the effects of the changes relative to EDM, three independent experiments were conducted using CD34+ cells from three mPBSC donors; HSPCs were cultured in EDM for phases 1 and 2 (for expansion and differentiation), and the culture was continued in EDM or EDM-TM-2 for phases 3 and 4 (**Figure 1A**). On average, a 6400-fold expansion was obtained on day 15 (**Figure 1B**). From days 15 to 32, 96%-99% of cells cultured with EDM or EDM-TM-2 expressed CD235a (**Figures 1C**), indicating full erythroid commitment. The loss of 98%-99% of CD36 expression confirmed efficient erythroid differentiation into reticulocytes with both approaches (**Figure S3A**).

Relative to EDM, EDM-TM-2 yielded significantly higher rates of enucleation on day 19 (mean: 94% vs. 76%) and day 22 (mean: 97% vs. 80%, all p <0.001; **Figure 1D**). In addition, EDM-TM-2 performed significantly better than EDM with respect to post-enucleation cell maintenance on days 22, 26, 28, and 32 (all p <0.05; **Figure 1E**).

Reticulocyte maturation was then characterized. Both approaches generally resulted in a similar loss of mitochondria (**Figure 1F**) and residual RNA (**Figure S3B**), showing that a certain degree of reticulocyte maturation had occurred with both approaches. Obtaining CD71-negative (CD71-) RBCs is the common confirmation that complete reticulocyte maturation is achieved. In this regard, EDM-TM-2 performed significantly better than EDM, as measured on days 22, 26, and 32 (all p<0.001; **Figure 1G**), confirming enhanced and near-complete reticulocyte maturation. Together, these observations show that EDM-TM-2 yields higher rates of terminal reticulocyte maturation than EDM, along with improved cell maintenance.

### EDM-TM-2 enhances cRBC survival during storage

Next, we tested whether the cRBCs produced with EDM-TM-2 could be stored in a nutritive solution commonly used by blood banks to store blood units. The mPBSCs of two donors were cultured with EDM-TM-2, and the resulting cRBCs were transferred to AS-3 solution (**Figure S4 A-E**). Through 42 days of storage at 4 °C (i.e., day 74 post-culture initiation), the number and proportion of total and CD235a+ cells (>90%) remained stable, confirming the persistence of EDM-TM-2-derived cRBCs (**Figure 2A and Figures S4I**). cRBCs maintained a normal appearance after 42 days of storage (as shown by GIEMSA staining [**Figure 2C**]) and some displayed a biconcave shape after 28 days of storage (as shown by confocal microscopy [**Figure S4K**]). Finally, serological test results of cRBCs (i.e., O+ and A+) were consistent with the blood types of mPBSC donors (**Figure 2B**). Crucially, the maturation phase of EDM-TM-2 was important for cell storage, since cells deprived of this phase (i.e. cultured up to the reticulocyte stage in EDM; **Figure S4 F-H**) tended to die more rapidly (all p <0.01 on days 28, 35, and 42 [**Figure 2A**]). Together, these results support that EDM-TM-2 produces mature cRBCs that can be preserved up to 42 days in storage solution without any adverse impact on phenotype.

## DISCUSSION

Building on other studies, we developed an *in vitro* RBC culture that yields erythroid proliferation, differentiation, and enucleation, with high levels of terminal maturation. The approach herein described does not use accessory or feeder cells, despite their acknowledged benefit for the proliferation, enucleation, and maturation processes.^13,16,17,47^

These conditions were chosen because accessory and feeder cells might interfere with the efficiency and characterization of gene editing, and because avoiding feeder cells is preferable for clinical applications. Despite this technical challenge, our results show that EDM-TM (designating both EDM-TM-1 and EDM-TM-2 except when specified otherwise) offers notable improvements over other existing methods.

First, EDM-TM enhanced the post-enucleation cell integrity of cRBCs — a key feature rarely reported in prior studies. On average, 48% of cells were maintained on day 32 — a 9-fold improvement compared to EDM. Giarratana et al. described EDM with feeder-free conditions in 5% plasma and reported a cell loss of ~40% following enucleation between days 18 and 24.^10^ We obtained similar results with EDM, however cell loss substantially increases further beyond that point in this medium. With another protocol developed by Heshusius et al., ~50% seemed to die after 12 days in differentiation medium,^13^ even though this protocol uses accessory cells. Unlike previous protocols, the present approach therefore enables the survival of cRBCs over an extended period of time post-enucleation, despite the lack of accessory and feeder cells.

Second, the levels of terminal maturation achieved with EDM-TM by day 32 (15%-20% of CD71+ cells) were higher than those reported in previous studies conducted without accessory cells (35%-80% of CD71+ cells).^7–12,14,32,34–36,48^ Interestingly, the level of terminal maturation achieved with EDM-TM was similar to those obtained in protocols that relied on accessory cells (10%-20% CD71+ cells)^13^ or feeder cells (5%-10% CD71+ cells).^16^

Third, EDM-TM-derived cRBCs could be preserved for as long as 42 days in an RBC nutritive solution used by blood banks, an improvement compared with the ≤28 days reported in prior studies.^10,14,33^ Some but not all cRBCs displayed a biconcave shape, which is expected in stored products.^45,46^ This practical benefit may be particularly important for clinical applications, since blood banks typically store blood units 42 days post-collection. The purpose of the EDM-TM method was further highlighted with the observation that EDM-derived immature reticulocytes were not as stable in these storage conditions as the EDM-TM-matured cRBC.

Consistent with previous studies,^7,13^ plasma had a key role in maintaining cell integrity. A 20% human plasma concentration was also necessary to achieve optimal levels of enucleation and cell maintenance in accessory-cell- and animal-serum-free conditions. This result is consistent with the poor enucleation rates and cell viability observed in serum-free cultures.^8,48^ Although an entirely synthetic medium may be preferable, human plasma therefore appears as an acceptable compromise for many applications, especially given the above-mentioned improvements of EDM-TM relative to other approaches.

Costs represent a major hurdle for the *in vitro* production of cRBCs,^49^ and EDM-TM-2 partially addresses this limitation. In EDM-TM-2, transferrin was either reduced from 1 mg/ml to 330 μg/ml on and after day 22; or replaced with FeIII-EDTA from days 12 to 22 (as tested by Olivier et al.).^31^ Of note, FeIII-EDTA could not be removed on and after day 22 without impairing maturation, showing a persistent role for iron during the later stages of differentiation. Moreover, while both SCF and EPO are essential for earlier stages of erythropoiesis, these reagents were dispensable later during the culture. Altogether, we estimate that these changes can reduce costs by ~600$ CAD per liter of medium, with virtually no changes in cRBC yields and characteristics.

In conclusion, EDM-TM provides a cell culture workflow compatible with both clinical applications and genome editing, in addition to providing a platform for studying terminal maturation mechanisms.

## ACKNOWLEDGEMENTS

This work was supported by a grant from MITACS and Héma-Québec. Y.B. holds a MITACS Accelerate graduate scholarship and was previously supported by a Canada graduates scholarship from CIHR. Salary support was provided by the Fonds de la Recherche du Québec-Santé (FRQS) to Y.D. The authors thank Samuel Rochette for feedback on the manuscript draft, Pascal Rouleau for expert technical help and Antoine Lewin for validating statistical analyses. We acknowledge the Bioimaging platform of the Infectious Disease Research Centre (CHU de Québec, Québec, QC, Canada), which is funded by an equipment and infrastructure grant from the Canadian Foundation Innovation (CFI).

## AUTHOR’S CONTRIBUTIONS

Conceptualization, Y.B., Y.D. and J.L.; methodology, Y.B, N.D. and J.L.; writing original draft, Y.B.; writing, review and editing, Y.B., Y.D. and J.L.; supervision, Y.D and J.L.; funding acquisition, Y.D and J.L. All authors reviewed the manuscript and approved its final version.

## REFERENCES

1. Chen K, Liu J, Heck S, Chasis JA, An X, Mohandas N. Resolving the distinct stages in erythroid differentiation based on dynamic changes in membrane protein expression during erythropoiesis. Proc Natl Acad Sci U S A. 2009;106(41):17413–17418.

2. Nandakumar SK, Ulirsch JC, Sankaran VG. Advances in understanding erythropoiesis: evolving perspectives. Br J Haematol. 2016;173(2):206–218.

3. Malleret B, Xu F, Mohandas N, et al. Significant biochemical, biophysical and metabolic diversity in circulating human cord blood reticulocytes. PLoS One. 2013;8(10):e76062.

4. Ovchynnikova E, Aglialoro F, von Lindern M, van den Akker E. The Shape Shifting Story of Reticulocyte Maturation. Front Physiol. 2018;9:829.

5. Liu J, Guo X, Mohandas N, Chasis JA, An X. Membrane remodeling during reticulocyte maturation. Blood. 2010;115(10):2021–2027.

6. Chasis JA, Prenant M, Leung A, Mohandas N. Membrane assembly and remodeling during reticulocyte maturation. Blood. 1989;74(3): 1112-1120.

7. Han SY, Lee EM, Choi HS, Chun BH, Baek EJ. The effects of plasma gelsolin on human erythroblast maturation for erythrocyte production. Stem Cell Res. 2018;29:64–75.

8. Kim SH, Lee E, Han SY, Choi HS, Ryu KY, Baek EJ. Improvement of red blood cell maturation in vitro by serum-free medium optimization. Tissue Eng Part C Methods. 2019.

9. Timmins NE, Athanasas S, Gunther M, Buntine P, Nielsen LK. Ultra-high-yield manufacture of red blood cells from hematopoietic stem cells. Tissue Eng Part C Methods. 2011;17(11):1131–1137.

10. Giarratana MC, Rouard H, Dumont A, et al. Proof of principle for transfusion of in vitro-generated red blood cells. Blood. 2011;118(19):5071–5079.

11. Baek EJ, Kim HS, Kim JH, Kim NJ, Kim HO. Stroma-free mass production of clinical-grade red blood cells (RBCs) by using poloxamer 188 as an RBC survival enhancer. Transfusion. 2009;49(11):2285–2295.

12. Leberbauer C, Boulme F, Unfried G, Huber J, Beug H, Mullner EW. Different steroids co-regulate long-term expansion versus terminal differentiation in primary human erythroid progenitors. Blood. 2005;105(1):85–94.

13. Heshusius S, Heideveld E, Burger P, et al. Large-scale in vitro production of red blood cells from human peripheral blood mononuclear cells. Blood Adv. 2019;3(21):3337–3350.

14. Zhang Y, Wang C, Wang L, et al. Large-Scale Ex Vivo Generation of Human Red Blood Cells from Cord Blood CD34(+) Cells. Stem Cells Transl Med. 2017;6(8):1698–1709.

15. Lee E, Sivalingam J, Lim ZR, et al. Review: In vitro generation of red blood cells for transfusion medicine: Progress, prospects and challenges. Biotechnol Adv. 2018;36(8):2118–2128.

16. Giarratana MC, Kobari L, Lapillonne H, et al. Ex vivo generation of fully mature human red blood cells from hematopoietic stem cells. Nat Biotechnol. 2005;23(1):69–74.

17. van den Akker E, Satchwell TJ, Pellegrin S, Daniels G, Toye AM. The majority of the in vitro erythroid expansion potential resides in CD34(-) cells, outweighing the contribution of CD34(+) cells and significantly increasing the erythroblast yield from peripheral blood samples. Haematologica. 2010;95(9):1594–1598.

18. Chu SH, Packer M, Rees H, et al. Rationally Designed Base Editors for Precise Editing of the Sickle Cell Disease Mutation. Crispr j. 2021;4(2):169–177.

19. Wu Y, Zeng J, Roscoe BP, et al. Highly efficient therapeutic gene editing of human hematopoietic stem cells. Nat Med. 2019.

20. Zeng J, Wu Y, Ren C, et al. Therapeutic base editing of human hematopoietic stem cells. Nature Medicine. 2020.

21. Canver MC, Smith EC, Sher F, et al. BCL11A enhancer dissection by Cas9-mediated in situ saturating mutagenesis. Nature. 2015;527(7577):192–197.

22. Magis W, DeWitt MA, Wyman SK, et al. High-level correction of the sickle mutation is amplified in vivo during erythroid differentiation. iScience. 2022:104374.

23. DeWitt MA, Magis W, Bray NL, et al. Selection-free genome editing of the sickle mutation in human adult hematopoietic stem/progenitor cells. Sci Transl Med. 2016;8(360):360ra134.

24. Hoban MD, Lumaquin D, Kuo CY, et al. CRISPR/Cas9-Mediated Correction of the Sickle Mutation in Human CD34+ cells. Mol Ther. 2016;24(9):1561–1569.

25. Park SH, Lee CM, Dever DP, et al. Highly efficient editing of the beta-globin gene in patient-derived hematopoietic stem and progenitor cells to treat sickle cell disease. Nucleic Acids Res. 2019.

26. Pattabhi S, Lotti SN, Berger MP, et al. In Vivo Outcome of Homology-Directed Repair at the HBB Gene in HSC Using Alternative Donor Template Delivery Methods. Mol Ther Nucleic Acids. 2019;17:277–288.

27. Christaki EE, Politou M, Antonelou M, et al. Ex vivo generation of transfusable red blood cells from various stem cell sources: A concise revisit of where we are now. Transfus Apher Sci. 2019;58(1): 108-112.

28. Kurita R, Suda N, Sudo K, et al. Establishment of immortalized human erythroid progenitor cell lines able to produce enucleated red blood cells. PLoS One. 2013;8(3):e59890.

29. Trakarnsanga K, Griffiths RE, Wilson MC, et al. An immortalized adult human erythroid line facilitates sustainable and scalable generation of functional red cells. Nat Commun. 2017;8:14750.

30. Kobari L, Yates F, Oudrhiri N, et al. Human induced pluripotent stem cells can reach complete terminal maturation: in vivo and in vitro evidence in the erythropoietic differentiation model. Haematologica. 2012;97(12):1795–1803.

31. Olivier EN, Zhang S, Yan Z, et al. PSC-RED and MNC-RED: Albumin-free and low-transferrin robust erythroid differentiation protocols to produce human enucleated red blood cells. Exp Hematol. 2019;75:31–52.e15.

32. Griffiths RE, Kupzig S, Cogan N, et al. Maturing reticulocytes internalize plasma membrane in glycophorin A-containing vesicles that fuse with autophagosomes before exocytosis. Blood. 2012;119(26):6296–6306.

33. Kim HO, Baek EJ. Red blood cell engineering in stroma and serum/plasma-free conditions and long term storage. Tissue Eng Part A. 2012;18(1-2): 117-126.

34. Lee E, Han SY, Choi HS, Chun B, Hwang B, Baek EJ. Red blood cell generation by three-dimensional aggregate cultivation of late erythroblasts. Tissue Eng Part A. 2015;21(3-4):817–828.

35. Liu Q, Luo L, Ren C, et al. The opposing roles of the mTOR signaling pathway in different phases of human umbilical cord blood-derived CD34(+) cell erythropoiesis. Stem Cells. 2020.

36. Miharada K, Hiroyama T, Sudo K, Nagasawa T, Nakamura Y. Efficient enucleation of erythroblasts differentiated in vitro from hematopoietic stem and progenitor cells. Nat Biotechnol. 2006;24(10): 1255-1256.

37. Simamura E, Arikawa T, Ikeda T, et al. Melanocortins contribute to sequential differentiation and enucleation of human erythroblasts via melanocortin receptors 1, 2 and 5. PLoS One. 2015;10(4):e0123232.

38. Ferrari G, Thrasher AJ, Aiuti A. Gene therapy using haematopoietic stem and progenitor cells. Nat Rev Genet. 2020.

39. Byrnes C, Lee YT, Meier ER, Rabel A, Sacks DB, Miller JL. Iron dose-dependent differentiation and enucleation of human erythroblasts in serum-free medium. J Tissue Eng Regen Med. 2016;10(2):E84–89.

40. Hu J, Liu J, Xue F, et al. Isolation and functional characterization of human erythroblasts at distinct stages: implications for understanding of normal and disordered erythropoiesis in vivo. Blood. 2013;121(16):3246–3253.

41. Boehm D, Murphy WG, Al-Rubeai M. The effect of mild agitation on in vitro erythroid development. J Immunol Methods. 2010;360(1-2):20–29.

42. Wickrema A, Krantz SB, Winkelmann JC, Bondurant MC. Differentiation and erythropoietin receptor gene expression in human erythroid progenitor cells. Blood. 1992;80(8):1940–1949.

43. Muta K, Krantz SB, Bondurant MC, Wickrema A. Distinct roles of erythropoietin, insulin-like growth factor I, and stem cell factor in the development of erythroid progenitor cells. J Clin Invest. 1994;94(1):34–43.

44. Poldee S, Metheetrairut C, Nugoolsuksiri S, Frayne J, Trakarnsanga K. Optimization of an erythroid culture system to reduce the cost of in vitro production of red blood cells. MethodsX. 2018;5:1626–1632.

45. Acker JP, Marks DC, Sheffield WP. Quality Assessment of Established and Emerging Blood Components for Transfusion. J Blood Transfus. 2016;2016:4860284.

46. Orlov D, Karkouti K. The pathophysiology and consequences of red blood cell storage. Anaesthesia. 2015;70 Suppl 1:29–37, e29-12.

47. Fujimi A, Matsunaga T, Kobune M, et al. Ex vivo large-scale generation of human red blood cells from cord blood CD34+ cells by co-culturing with macrophages. Int J Hematol. 2008;87(4):339–350.

48. Uchida N, Demirci S, Haro-Mora JJ, et al. Serum-free Erythroid Differentiation for Efficient Genetic Modification and High-Level Adult Hemoglobin Production. Mol Ther Methods Clin Dev. 2018;9:247–256.

49. Rousseau GF, Giarratana MC, Douay L. Large-scale production of red blood cells from stem cells: what are the technical challenges ahead? Biotechnol J. 2014;9(1):28–38.

