## Supplementary material for "Accessory-cell-free differentiation of hematopoietic stem and progenitor cells into mature red blood cells": Supp Material

#### SUPPLEMENTARY TABLES

##### Table S1: HSPC donors' characteristics

| Identification | Figure | Gender | Age | Ethnicity |
|---|---|---|---|---|
| **mPBSC #6** | 1 | M | 24 | Black |
| **mPBSC #7** | 1 | M | 29 | Black |
| **mPBSC #8** | 1 | M | 22 | Black |
| **mPBSC #9** | 2<br>S1<br>S2 | M | 30 | Other |
| **mPBSC #10** | 2<br>S1 | M | 31 | Black |
| **mPBSC #11** | 2<br>S1 | M | 26 | Other |
| **mPBSC #2** | S1 | M | 31 | White |
| **mPBSC #12** | S2 | M | 22 | Asian |
| **mPBSC #14** | 2 | M | 36 | White |
| **mPBSC #1** | S1 | M | 30 | Black |

#### Table S2: Catalog numbers and vendors

| Name | Vendor | | #catalog |
|---|---|---|---|
| **Purified mobilized peripheral blood stem cells** | AllCells | Alameda, CA, USA | mPB015F or mPB014F |
| **Iscove's Modified Dulbecco's Medium** | Sigma-Aldrich | Oakville, ON, Canada | 12440-079 |
| **Glutamine** | ThermoFisher | Ottawa, ON, Canada | 25030081 |
| **Insulin** | Sigma-Aldrich | Oakville, ON, Canada | I3536 |
| **Plasma**<br><br>**AB Octoplasma** | Héma-Québec | Québec, QC, Canada | - |
| | Octopharma | Toronto, ON, Canada | DIN02270013 |
| **Hydrocortisone** | Sigma-Aldrich | Oakville, ON, Canada | H0888 |
| **IL-3** | StemCell Technologies | Vancouver, BC, Canada | 78040 |
| **SCF** | R&D system | Minneapolis, MN, USA | 255-SC-200/CF |
| **EPO** | Peprotech | East Windsor, NJ, USA | 100-64 |
| **Transferrin** | R&D system | Minneapolis, MN, USA | 2914-HT |
| **FeIII-EDTA** | Sigma-Aldrich | Oakville, ON, Canada | E6760 |

| | | | |
|---|---|---|---|
| **Acrodisc WBC Syringe Filter 25 mm** | VWR | Mont-Royal, QC, Canada | CA10127-206 |
| **anti-CD71-APC** | BD Biosciences | Franklin Lakes, NJ, USA | 551374 |
| **anti-CD235a** | BD Biosciences | Franklin Lakes, NJ, USA | 559943 |
| **anti-CD47-BV421** | BD Biosciences | Franklin Lakes, NJ, USA | 563760 |
| **anti-CD36-FITC** | BD Biosciences | Franklin Lakes, NJ, USA | 555454 |
| **7AAD** | Beckman Coulter | Indianapolis, IN, USA | A07704 |
| **SYTO 13 green** | ThermoFisher | Ottawa, ON, Canada | S7575 |
| **MitoTracker™ Orange CMTMRos** | ThermoFisher | Ottawa, ON, Canada | M7510 |
| **RetiCount** | BD Biosciences | Franklin Lakes, NJ, USA | 349204 |
| **anti-AB** | BioRad | Saint-Laurent, QC, Canada | 801375 |
| **anti-A** | BioRad | Saint-Laurent, QC, Canada | 801325 |
| **anti-B** | BioRad | Saint-Laurent, QC, Canada | 801350 |
| **anti-D** | BioRad | Saint-Laurent, QC, Canada | 802033 |

### SUPPLEMENTARY FIGURES

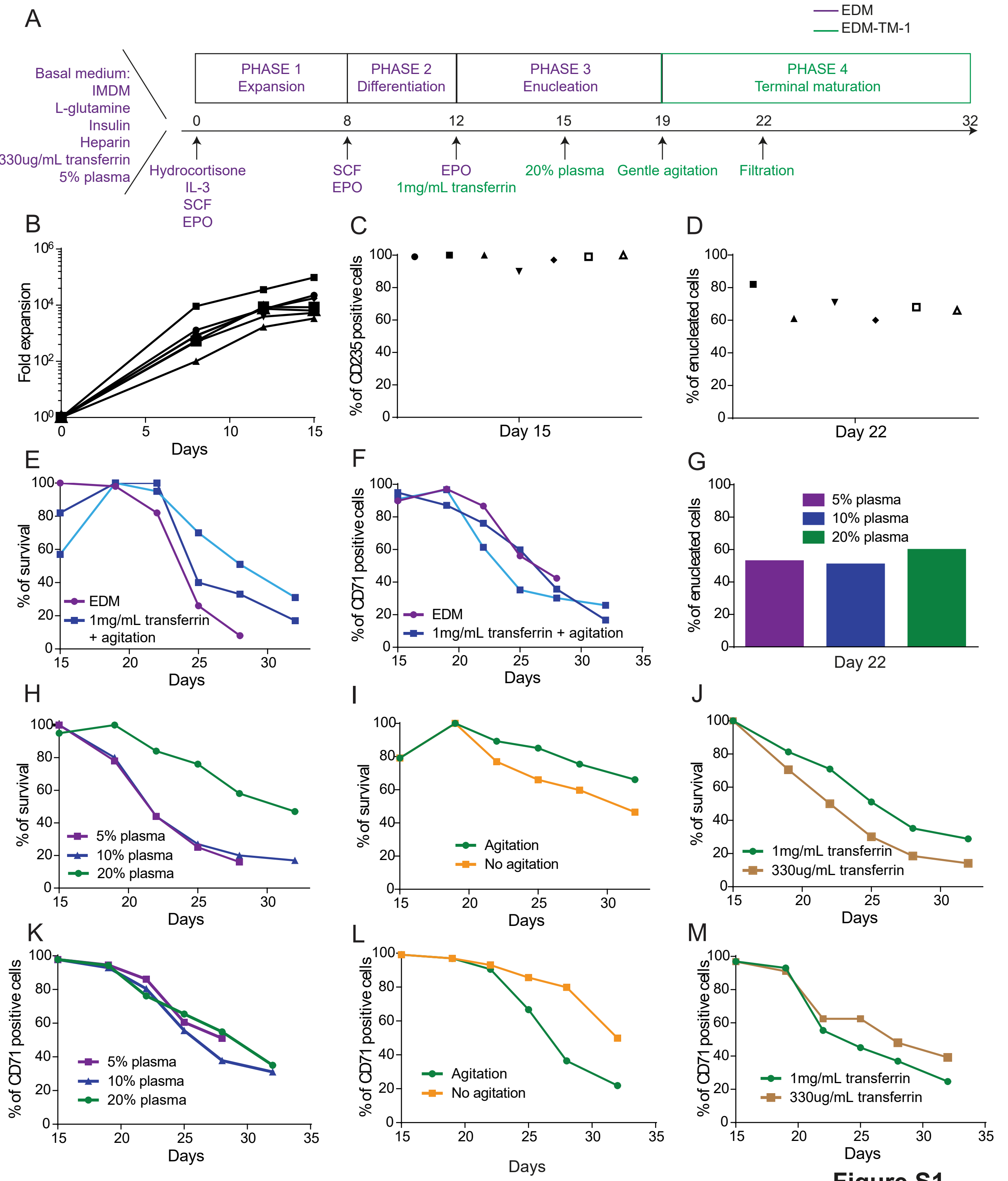


**Figure S1**
**Boccacci et al.**

**Figure S1: Development of EDM-TM-1: assessing the effect of individual changes.**

(A) Overview of EDM-TM-1, with modifications relative to EDM indicated in green.

(B-M) RBC survival is expressed as the percentage of the maximum number of cells obtained through the culture and RBC maturation is expressed as the decreasing percentage of CD71+ cells. EDM-TM-1 conditions are represented in green in the graphs, and EDM in purple. (B) Erythroid expansion of HSPCs from day 0 to day 15, (C) Percentage of CD235a positive cells on day 15, and (D) Percentage of enucleated cells on day 22, from all seven cultures used for optimization (six distinct donors); (E) Percentage of cRBC survival in EDM (●) or EDM supplemented with high transferrin levels from day 12 and mild agitation from day 19 from two distinct donors (■); (F) RBC maturation in EDM or EDM supplemented with high transferrin levels from day 12 and mild agitation from day 19 from two distinct donors; (G) Percentage of enucleated cRBCs on day 22 in different plasma concentrations; (H) Percentage of cRBC survival in different plasma concentrations; (I) Percentage of cRBC survival with (●) or without (■) agitation starting on day 19; (J) Percentage of cRBC survival with low (■) or high (●) transferrin levels starting on day 12; (K) RBC maturation, when cultured in different plasma concentrations; (L) RBC maturation, when cultured with high versus low transferrin levels starting on day 12; (M) RBC maturation, with or without agitation starting on day 19.

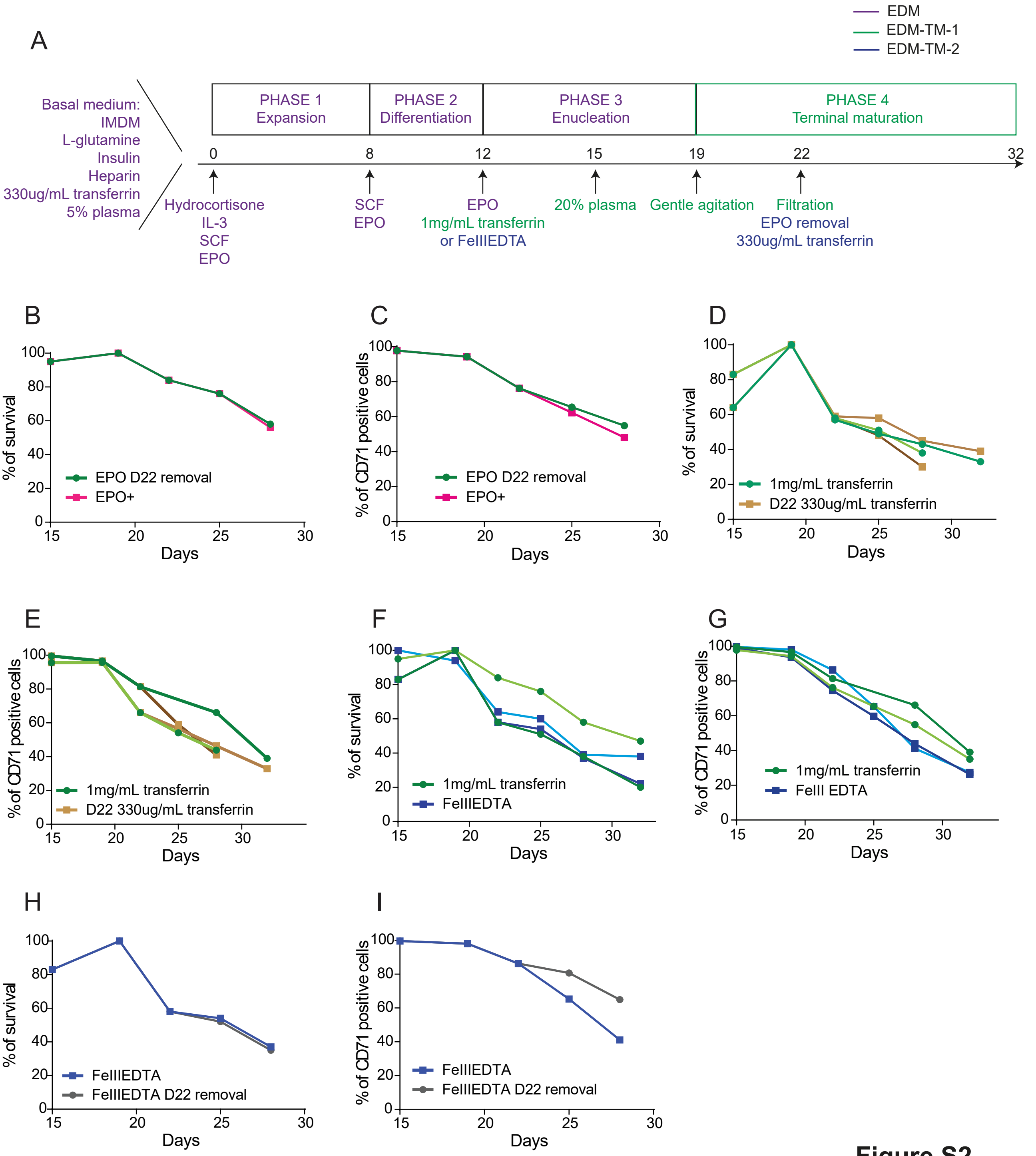


**Figure S2**
**Boccacci et al.**

**Figure S2: Development of EDM-TM-2 to reduce costs.** (A) Overview of EDM-TM-2, with modifications relative to EDM indicated in green and relative to EDM-TM-1 indicated in blue. (B-I) HSPCs from distinct donors are illustrated with different shades of the same color. RBC survival is expressed as the percentage of the maximum number of cells obtained through the culture and RBC maturation is expressed as the decreasing percentage of CD71+ cells. (B) Percentage of cRBC survival withdrawing (●) or keeping (■) EPO from day 22; (C) RBC maturation, withdrawing or keeping EPO from day 22. (D) Percentage of cRBC survival when cultured with a high transferrin concentration added from day 12 up until the end of the culture (●) or from days 12 to 22 (■); (E) RBC maturation, when cultured with a high transferrin concentration from day 12 up until the end of the culture, or from days 12 to 22. (F) Percentage of cRBC survival when cultured with high transferrin concentrations from day 12 (●) versus low transferrin concentration and 4 µM FeIII-EDTA (■) starting on day 12; (G) RBC maturation, when cultured with high transferrin concentrations from day 12 versus low transferrin concentration and 4 µM FeIII-EDTA. (H) Percentage of cRBC survival expressed when cultured with FeIII-EDTA up until end of cell culture (■) or until day 22 (●); (I) RBC maturation, when cultured with FeIII-EDTA up until end of cell culture versus up until day 22.

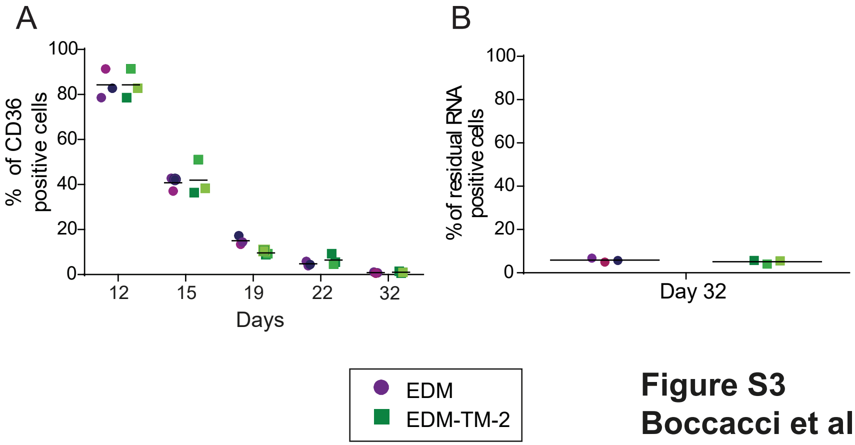


**Figure S3**
**Boccacci et al.**

**Figure S3: Differentiation and terminal maturation of EDM-TM-2 versus EDM-derived cRBCs**. (A) Percentage of CD36+ (early erythroid differentiation marker and expected to disappear at the reticulocyte stage) cells; (B) percentage of RetiCount+ (residual RNA marker) cells.*P <0.05 **P <0.01; ***P <0.001.

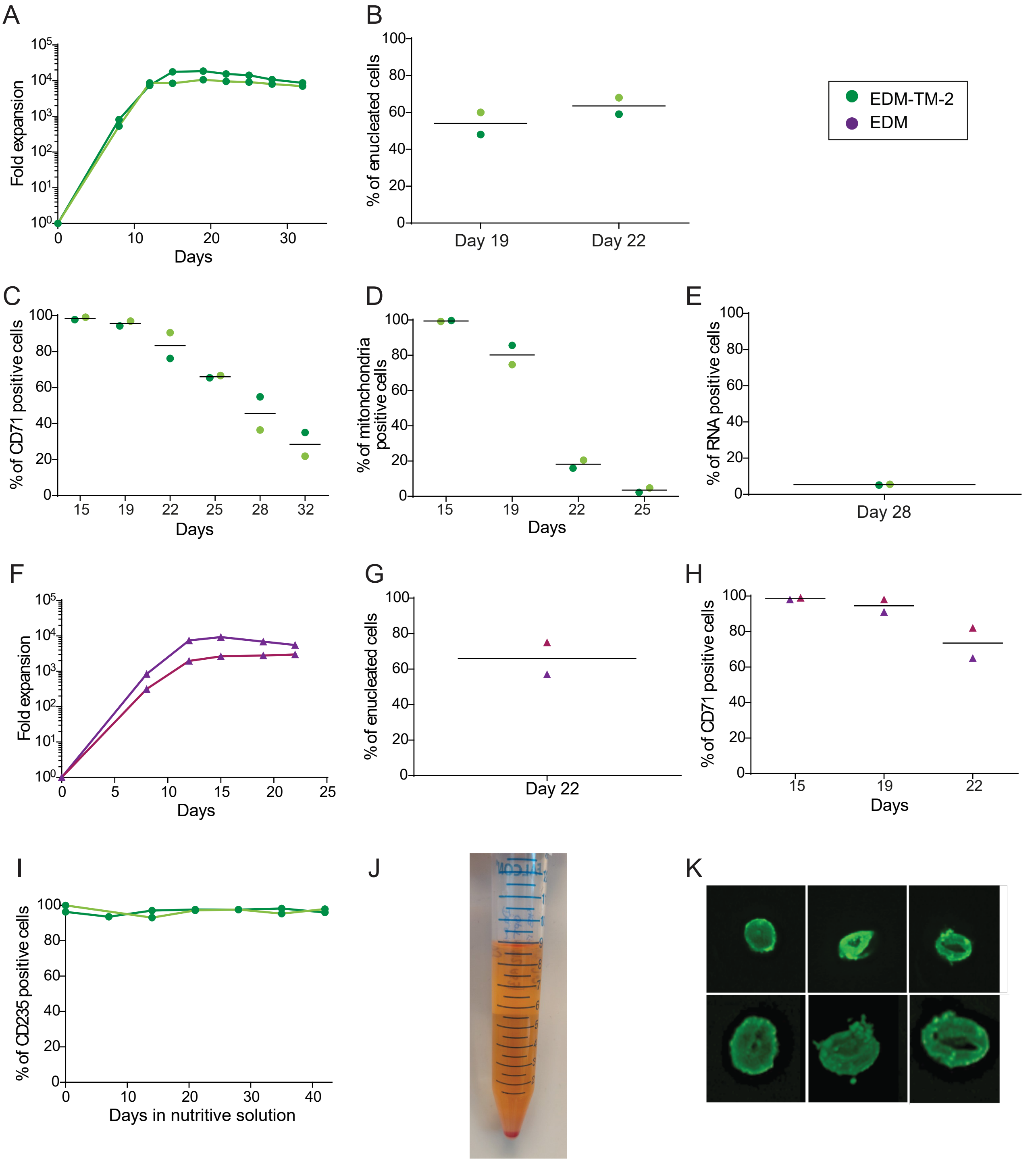


**Figure S4**
**Boccacci et al.**

**Figure S4: Characterization of RBCs cultured in EDM or EDM-TM-2 prior to and/or prior and during storage in nutritive solution.** HSPC-derived RBCs from distinct donors are represented with different shades of the same color.

(A) Erythroid expansion of HSPCs and maintenance of cRBCs with EDM-TM from day 0 to day 32; (B) Percentage of enucleated cells with EDM-TM (≥100 cells enumerated following GIEMSA staining); (C) Percentage of CD71+ (maturation marker) cells; (D) Percentage of MitoTracker+ (mitochondria marker) cells; (E) Percentage of RetiCount+ (residual RNA marker) cells. (F) Erythroid expansion of HSPCs with EDM from day 0 to day 22; (G) Enucleated, SYTO-Green+ cells on day 22 with EDM; (H) Percentage of CD71+ (maturation marker) cells. (I) Percentage of CD235a+ cells (late erythroid differentiation marker and RBC marker) throughout storage of EDM-TM cRBCs; (J) Photo of pelleted RBCs stored at 4°C; (K) 3D reconstructions of the biconcave shape of EDM-TM cRBCs on day 28 of storage (day 60 post-protocol initiation) taken with confocal microscopy.
